## Supplementary material for "Dual human lung models reveal compartment-specific activity of anti-tuberculosis drugs and host-directed therapies"

### 1. SUPPLEMENTAL FIGURE LEGENDS

**Figure S1. Molecular characterization of AML and MDM and flow cytometry gating strategy for phenotypic analysis. A)** Relative gene expression in AML and MDM cells assessed by RT-qPCR. **B)** Immunoblot analysis of PPARγ expression in AML and MDM cells from four independent donors. Quantification of protein levels is shown. **C)** Flow cytometry gating strategy used for the analysis of hAMs. In the graphs, each dot represents an individual donor. Significant differences were determined using a paired, non-parametric Wilcoxon test. $p < 0.05$ (*), $p < 0.01$ (**), $p < 0.001$ (***).

**Figure S2. Differential gene expression analysis of AML versus MDM. A)** Heatmap of the principal representative genes. The heatmap shows differences in expression across the samples. Rows correspond to the different genes and columns to the different samples of AML and MDM cells. Red color is indicative of up-regulated genes, and blue is down-regulated; AdjPvalue < 0.05 is shown with *. **B)** Volcano plot shows the top of principal 40 DEG (FC +/- 1.5 and AdjPvalue < 0.05). The blue dots denote down-regulated genes, and the red dots denote up-regulated genes in AML. Grey dots are out of range. **C)** Gene Ontology (GO) enrichment analysis of biological processes (BP) for up-regulated genes with the principal deregulated genes between AML vs MDM cells. The dot size indicates the number of genes, and the colors correspond to the Adjusted P-Value.

**Figure S3. Optimization of infection conditions and phenotypic profiling of AML cells upon Mtb infection. A)** Uninfected and H37Rv-GFP-infected AML were visualized using epifluorescence and light microscopy. Bacteria are shown in green. Scale bars: 25 µm. **(B-C)** Uninfected and H37Rv-GFP-infected AML (MOI of 5:1, 1:1, 0.5:1, and 0.1:1) were analyzed by flow cytometry. The percentage of live cells (B) and GFP+ cells among live cells (C) is shown. **D)** Measurement of bacteria proliferation in infected AML (MOI 1:1) after 0-, 1-, 3-, or 6-days post-infection (p.i.) by using H37Rv::lux luminescent bacteria (left), colony-forming unit (CFU, center), or Median Fluorescence Intensity (MFI, right) of H37Rv-GFP strain. **E)** Median Fluorescence Intensity (MFI) of CD86, CD163, CD169, CD209, and MARCO of $CD16^{+}CD14^{+}$ and $CD16^{-}CD14^{+}$ subsets in uninfected and H37Rv-infected AML cells. Significant differences were determined using an unpaired, non-parametric Mann–Whitney test. $p < 0.05$ (*), $p < 0.01$ (**), $p < 0.001$ (***).

**Figure S4. Evaluation of toxicity induced by Anti-TB Agents in AML cells.**

Lactate dehydrogenase (LDH) release was measured as a marker of cytotoxicity. Mtb-infected AMLs (MOI 1:1) were treated with different compounds and incubated for 6 days.

**Supplementary Movie 1:** Ciliary beating of airway ALI culture visualized using light microscopy (EVOS M7000 - Invitrogen) with a 60× objective. Real-time recording (live) showing coordinated cilia movement.

### 2. SUPPLEMENTARY TABLES

**Table S1. List of commercial drugs used.**

| Name of the molecule | Mechanism relevant to TB control | Compound class |
|---|---|---|
| Isoniazid (INH) | Inhibits mycolic-acid synthesis (81, 82) | Standard-of-Care (SoC) |
| Moxifloxacin (MXF) | A fourth-generation fluoroquinolone antibiotic that inhibits DNA replication in Mtb by diminishing DNA topoisomerase II and IV activity (83, 84) | |
| Pyrazinamide (PZA) | A first-line agent whose activity and mechanism are still under investigation, with current evidence suggesting roles in disrupting membrane transport and proton motive force (85, 86) as well as other targets (87) | |
| Rifampicin (RIF) | A first-line ansamycins antibiotic that specifically binds to the β subunit of bacterial DNA-dependent RNA polymerase, thereby inhibiting transcription (88, 89) | |
| Linezolid (LZD) | An oxazolidinone antibiotic that inhibits bacterial protein synthesis by binding to the 50S ribosomal subunit (90, 91) | |
| Ibuprofen (IBU) | A non-steroidal anti-inflammatory drug (NSAID) that inhibits COX-1/2 and reduces $PGE_2$ synthesis, enhancing phagocytosis, nitric oxide (NO) production, and NADPH oxidase activity (53, 92) | Host-Directed Therapies (HDT) |
| Aspirin (ASA) | A NSAID with antiplatelet, anti-inflammatory, and analgesic properties that inhibits COX-1/2 and reduces $PGE_2$ synthesis; additionally, it promotes the formation of aspirin-triggered lipoxins that help resolve inflammation (92, 93) | |
| Doramapimod (DORA) | Selective inhibitor of p38 MAPK, reducing pro-inflammatory cytokine production (52, 94) | |
| Simvastatin (SMV) | Enhances phagosome maturation and promotes apoptosis and autophagy in Mtb-infected cells (95–97) | |
| Metformin (MET) | Induces reactive oxygen species (ROS) production and increases phagosome–lysosome fusion (98) | |
| Ethoxzolamide (ETZ) | A carbonic anhydrase inhibitor that has been repurposed for TB research by targeting the PhoPR two-component regulatory system in Mtb (99) | Virulence-Targeting Therapies (VTT) |
| BBH7 | Inhibits the ESX-1 secretion system, impairing virulence factor export (76, 100) | |

**Table S2. List of SYBR Green human primers used for qPCR.**

| Primers | Gene reference | Sequences - Forward (top), Reverse (bottom) |
|---|---|---|
| *MRC1* | NM_002438.4 | GCCTCGTTGTTTTGCGTCTT |
| | | GAGAACAGCACCCGGAATGA |
| *MARCO* | NM_006770.4 | AGGAGGACGAGCTCTTGAGT |
| | | TCAGAACTTGGACCACCAGC |
| *CXCL3* | NM_002090.3 | CCCAAACCGAAGTCATAGCCA |
| | | ACCCTGCAGGAAGTGTCAAT |
| *PPARG* | NM_138711.6 | GGTGACCAGAAGCCTGCATT |
| | | CACGGAGCTGATCCCAAAGT |
| *DUSP1* | NM_004417.4 | GTACTAGCGTCCCTGACAGC |
| | | CCCAGGTACAGAAAGGGCAG |
| *SPI1* | NM_001080547.2 | AAAATCAGGAACTTGTGCTGGC |
| | | GGGGAAACCCTTCCATTTTGC |
| *TNF* | NM_000594.4 | GAGGCCAAGCCCTGGTATG |
| | | CGGGCCGATTGATCTCAGC |
| *IL10* | NM_000572.3 | ACTTTAAGGGTTACCTGGGTTGC |
| | | TCACATGCGCCTTGATGTCTG |
| *MMP7* | NM_002423.5 | GTCTCTGGACGGCAGCTATG |
| | | TAGTCCTGAGCCTGTTCCCA |
| *MMP9* | NM_004994.3 | GGACAAGCTCTTCGGCTTCT |
| | | TCGCTGGTACAGGTCGAGTA |
| *IFNA* | NM_024013.3 | GTGAGGAAATACTTCCAAAGAATCAC |
| | | TCTCATGATTTCTGCTCTGACAA |
| *IFNB* | NM_002176.4 | AGCTGCAGCAGTTCCAGAAG |
| | | AGTCTCATTCCAGCCAGTGC |
| *MX1* | NM_001144925.2 | CGGAATCTTGACGAAGCCTG |
| | | CCTTTCCTTCCTCCAGCAGA |
| *MX2* | NM_002463.2 | CAGAGGCAGCAGACGATCAAC |
| | | TTGGTCAGGATACCGATGGTC |
| *ISG15* | NM_005101.4 | CGCAGATCACCCAGAAGATCG |
| | | TTCGTCGCATTTGTCCACCA |
| *SIGLEC1* | NM_023068.4 | ATGGGGTACGCCTCCAAAC |
| | | GTGCCTCATTGGGTGTGTTG |
| *YWHAZ* | NM_001135699.2 | CCTGCATGAAGTCTGTAACTGAG |
| | | GACCTACGGGCTCCTACAACA |

**Table S3. List of antibodies used for Western Blotting.**

| Antibody (species) | Dilution | Supplier | Reference |
|---|---|---|---|
| PPARγ (rabbit) | 1:1000 | Cell Signaling Technology | 2435 |
| β-actin (mouse) | 1:1000 | Merck | A1978 |
| Anti-rabbit (goat) | 1:5000 | Advansta | R-05072-500 |
| Anti-mouse (goat) | 1:5000 | Advansta | R-05071-500 |

**Table S4. List of antibodies used for spectral Flow Cytometry.**

| Antibody | Fluorochrome | Clone | Supplier | Catalog # | Dilution |
|---|---|---|---|---|---|
| ViaDye | Red | - | Cytek | R7-60008 | 1:4000 |
| CD206 | AF700 | 15-2 | Biolegend | 321132 | 1:400 |
| CD14 | CfluorB548 | 63D3 | Cytek | R7-20116 | 1:200 |
| CD169 | BV421 | 7-239 | Biolegend | 346018 | 1:100 |
| CD4 | BV650 | SK3 | Cytek | R7-20166 | 1:100 |
| CXCR4 | PE-Cy5 | 12G5 | Biolegend | 306507 | 1:50 |
| CCR5 | BV785 | J418F1 | Biolegend | 359131 | 1:50 |
| CD209 | APC fire750 | 9E9A8 | Biolegend | 330115 | 1:100 |
| CD86 | BV605 | IT2,2 | Biolegend | 305429 | 1:50 |
| PDL1 | PE-fire810 | 29E,2A3 | Biolegend | 329755 | 1:100 |
| CD64 | BV650 | 10,1 | Biolegend | 305053 | 1:50 |
| CD163 | PE-Dazzle594 | GHI/61 | Biolegend | 333623 | 1:100 |
| MertK | BV711 | 590H11G1E3 | Biolegend | 367619 | 1:200 |
| CD16 | CfluorV450 | 3G8 | Cytek | R7-20184 | 1:50 |
| MARCO | PE-Cy7 | PLK-1 | Thermofisher | 25-5447-42 | 1:20 |
| CD36 | PE | CB38(NL07) | Thermofisher | A15793 | 1:20 |
| ABCA1 | Dylight 650 | polyclonal | Thermofisher | PA5-22907 | 1:20 |
| CD45 | BV510 | HI30 | Biolegend | 304035 | 1:50 |
| CD38 | PE-Fire640 | S17015F | Biolegend | 397217 | 1:50 |
