## Supplementary figures and images for "Dual human lung models reveal compartment-specific activity of anti-tuberculosis drugs and host-directed therapies"

### Supp. Figure 1

#### Supp. Figure 1

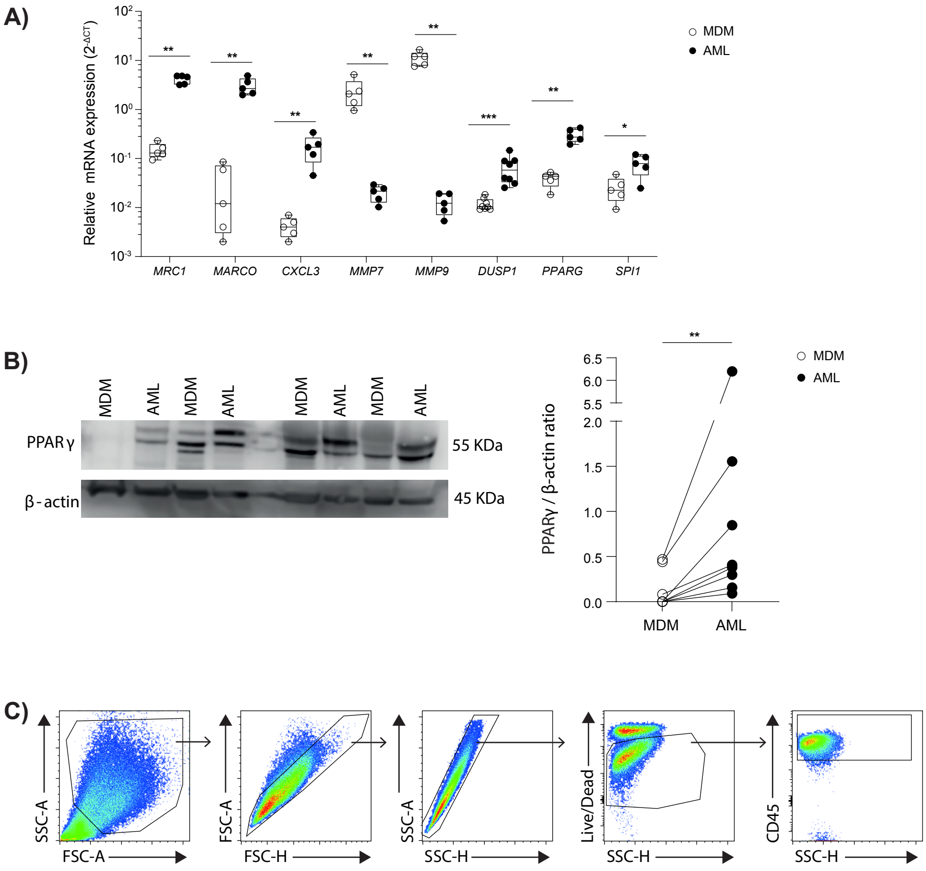

### Supp. Figure 2

#### Supp. Figure 2

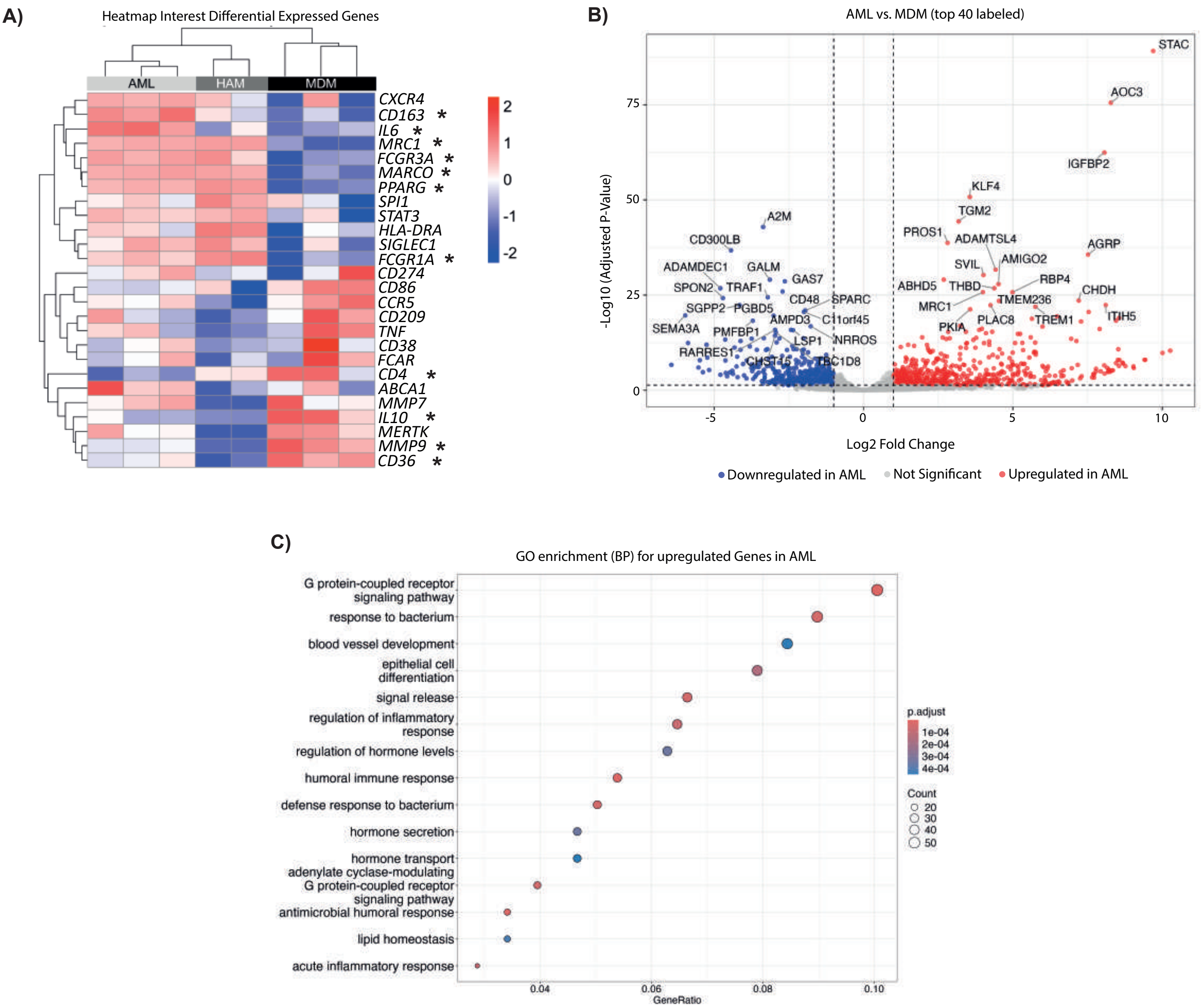

### Supp. Figure 3

#### Supp. Figure 3

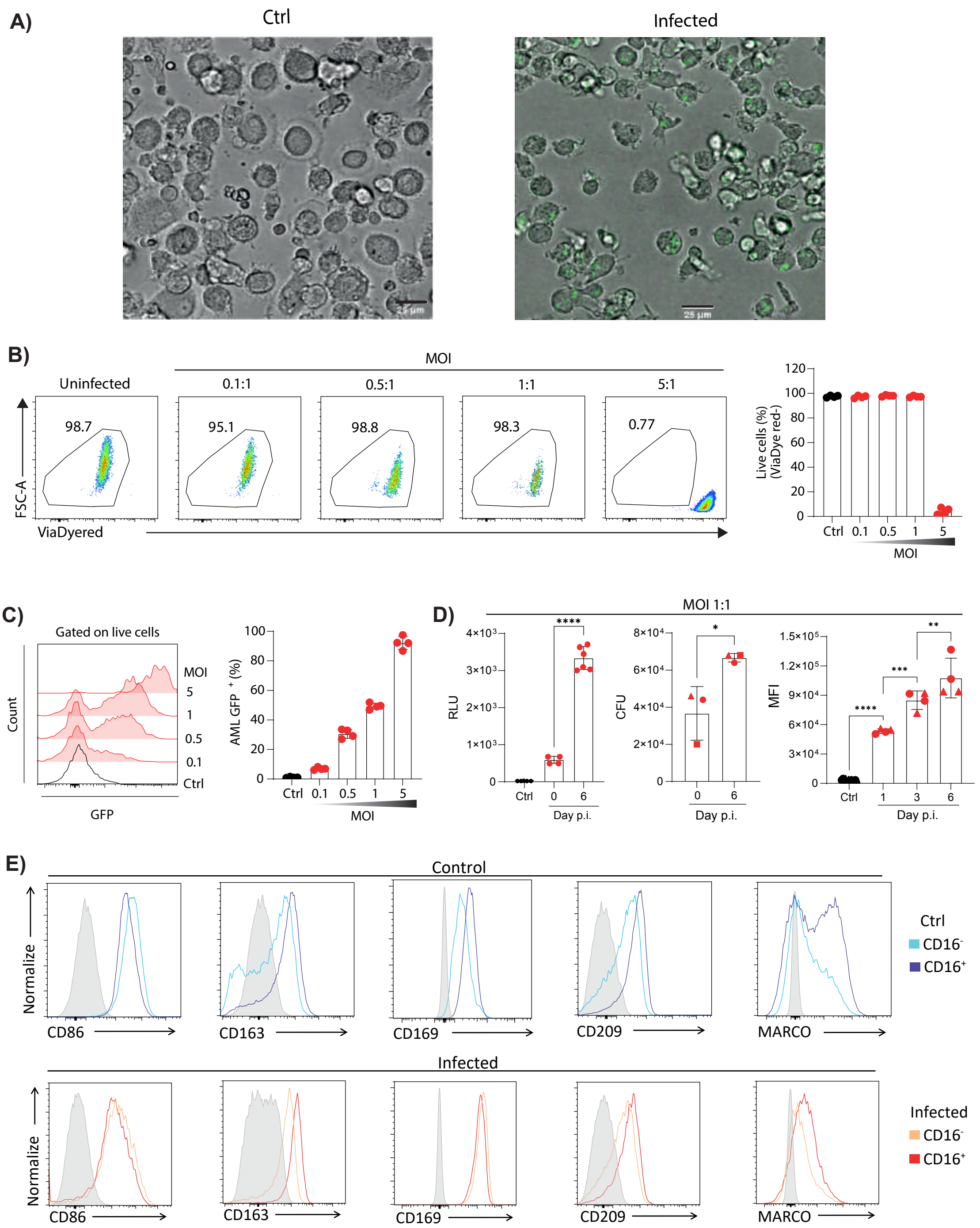

### Supp. Figure 4

Supp. Figure 4

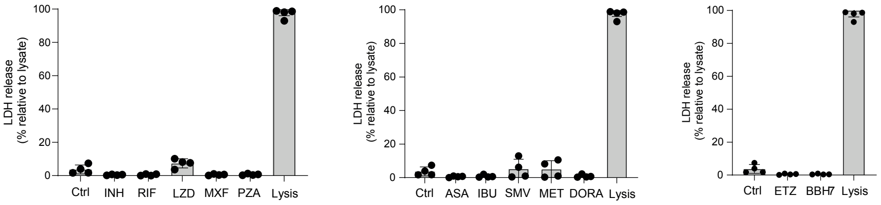
